# Cerebrospinal fluid transcriptional analyses reveals upregulation of IL-17, Type 1 interferon transcriptional pathways and neutrophil persistence genes associated with increased mortality from pneumococcal meningitis in adults

**DOI:** 10.1101/490045

**Authors:** Emma C Wall, José Afonso Guerra-Assunção, Gabriele Pollara, Cristina Venturini, Veronica S Mlozowa, Theresa J Allain, David G Lalloo, Mahdad Noursadeghi, Jeremy S Brown, Robert S Heyderman

## Abstract

**Background:** Improving outcomes from pneumococcal meningitis (PM), particularly in populations with high HIV prevalence, requires better understanding of host inflammatory responses to infection.

**Methods:** We compared the transcriptome in pre-antibiotic cerebrospinal fluid (CSF) and blood from Malawian adults with PM using RNA sequencing. We used network analyses and cellular/process deconvolution of the transcriptome to identify important patho-physiological associations with outcome.

**Findings:** Blood transcriptional profiles were obtained in 28 patients (21 HIV co-infected; median age 33 years [26-66]; median CSF WCC 28 cells/mm^3^ [0-3660]; median bacterial load 4.7×10^6^ copies/ml CSF [671-2×10^9^]; in-hospital mortality 64%), paired CSF profiles were obtained in 13. Marked differences in gene expression by outcome were confined to the CSF. In non-survivors, differentially expressed genes in the CSF were co-correlated in a network of pro-inflammatory gene-clusters enriched for collagen degradation and platelet degranulation. In contrast, CSF gene expression networks from surviving patients were dominated by DNA repair, transcriptional regulation and immunological signalling. CSF expression of gene response-modules for IL-17, Type 1 interferons and IL-10 were enriched in non-survivors, expression of cell-specific response-modules did not differ by outcome. However, genes for neutrophil chemotaxis and persistence were highly over-expressed in non-survivors.

**Interpretation:** These data suggest poor outcome in PM is associated with over-expression of IL-17 and T1-IFN associated pro-inflammatory responses in the CSF and suggest a role for neutrophil-mediated inflammation. These responses are unlikely to be effected by current adjunctive treatments. Improving poor outcomes from PM will require better-targeted interventions.

**Funding:** Academy of Medical Sciences (UK), Wellcome Trust (UK) (089671/B/09/Z)

## Background

*Streptococcus pneumoniae* remains the most prevalent cause of community acquired meningitis-associated mortality and morbidity in adults in most settings.^1-3^ The greatest burden of pneumococcal meningitis (PM) falls in low and middle income countries (LMICs) with high HIV prevalence. Estimates of incidence in adults and adolescents in these settings varies between 3-40/100 000, compared to 0.2- 1.5/100 000 in high resource, low HIV prevalence settings.^4-8^ In LMICs with high HIV prevalence, the reported mortality in adults and adolescents from PM is 50-70%.^9,10^

Animal model and human post-mortem studies indicate that disease pathogenesis in PM is characterised by a marked inflammatory response to bacterial invasion in the CSF, rather than an inability to mount an immune response. The inflammatory cascade and cytotoxic effects of host pro-inflammatory mediators^11,12^ together with bacterial toxins drive tissue damage characterised by apoptotic neuronal cell injury, raised intracranial pressure (ICP), thrombosis, cerebral oedema, and cerebral ischaemia.^13,14^ These findings supported the use of anti-inflammatory agents, such as dexamethasone as adjunct therapies for PM. Dexamethasone has demonstrated efficacy in controlled trials in industrialised countries in HIV-negative adults with PM, with an estimated relative risk reduction in mortality of 0.5 (95% CI 0.3 – 0.83)^15^. However, adjunctive dexamethasone and glycerol therapies have proven to be ineffective or even harmful in LMIC settings when tested in controlled trials, irrespective of HIV-1 serostatus.^2,16,17^ The differences in in overall case fatality rates by geographical location and response to dexamethasone in patients with PM are unexplained. Patients with bacterial meningitis in LMICs are younger, have a higher incidence of HIV co-infection, present later, and have important differences in bedside predictors of poor outcome compared to patients in better resourced settings.^18,19^ HIV-1 co-infection is not associated with worse outcomes from PM in any setting.^18-20^ Data from animal models and from patients from industrialised countries suggest excessive inflammatory responses determine disease severity, but this may not reflect the situation in LMICs.

Disease-specific immunomodulatory transcriptomic signatures have been identified that are strongly associated with disease severity and outcome in sepsis and influenza.^21-23^ In contrast, published reports on the transcriptomic response in bacterial meningitis are limited to human brain-endothelial cell lines and animal models,^24,25^ data in human disease are lacking

Based on our previously reported observation of a strong association between poor outcome and low CSF leukocyte counts in patients recruited to our centre,^19^ we tested the hypothesis that neutrophil-specific transcriptional activity would be down-regulated in both blood and CSF in non-survivors of PM compared to survivors. We report the results of our investigation into the blood and CSF host transcriptomic responses in Malawian patients with *S. pneumoniae* meningitis on admission to hospital. We have used network analyses and cellular and process deconvolution of the blood and CSF transcriptome to identify important components of the host inflammatory response associated with poor outcomes from pneumococcal meningitis.

## Methods

### Participants

Adults and adolescents presenting to Queen Elizabeth Central Hospital in Blantyre, Malawi with proven bacterial meningitis caused by *S. pneumoniae* between 2011-2013 were included (Current Controlled Trials registration ISRCTN96218197)^2^. All CSF and blood samples were collected prior to administration of parenteral ceftriaxone 2g BD for 10 days.^16,17^ Clinical data are from the first recording on admission to hospital, follow up was done to six weeks post-discharge.^2^

### Procedures

Routine CSF microscopy, cell count, and CSF culture was done at the Malawi-Liverpool-Wellcome Trust Clinical Research Programme laboratory in Blantyre, Malawi as previously described.^2^ Culture negatives samples were screened using the multiplex real-time polymerase chain reaction for *S. pneumoniae, N. meningitidis* and *Haemophilus influenzae type b* (Hib) kit from Fast-Track Diagnostics (FTD Luxemburg) according to the manufacturer’s instructions, bacterial loads were estimated from Ct values. We collected 2.5 ml of CSF and whole blood for transcriptional profiling in blood PAX-gene^®^ (Pre-AnalytiX, Qiagen, USA) tubes, incubated for 4 hours at room temperature, and stored at -80 degrees Celsius. Inhospital HIV testing was done on all patients by the clinical teams using point-of care Genie™ HIV1&2 test kits (BioRad, USA).

RNA was extracted from blood and CSF using the PAXgene^®^ Blood miRNA kit (Pre-Analytix, Qiagen, USA) according to the manufacturer’s instructions, with an additional mechanical disruption step in the CSF samples to disrupt the pneumococcal cell wall at 6200 rpm for 45 seconds in the Precellys evolution tissue homogenizer (Bertin Instruments). The extracted RNA was quantified and RNA Integrity Number (RIN) scores calculated using RNA Tapestation 4200^®^ (Agilent, USA) and Nanodrop^®^ (Thermoscientific, USA). Extracted RNA samples underwent library preparation for polyA tailed mRNA with a RNA concentration of >1ng/1ul using with Kapa RNA hyperPrep kit (Roche), followed by 75 cycles of Next-generation sequencing with NextSeq^®^ (Illumina, USA) by the Pathogen Genomics Laboratory at University College London.

### Bioinformatics and statistical analysis

All conventional statistical tests were two tailed, alpha <0.05 determined statistical significance. 95% confidence intervals are presented for odds ratios. Logistic regression was used to model associations between clinical outcomes and risk factors while controlling for confounding factors.

Sequence data quality was assessed prior to mapping by using the FASTQC toolkit (http://www.bioinformatics.babraham.ac.uk/projects/fastqc/). Mapping quality and percentage of properly mapped pairs, assessed from the BAM files, were considered for quality control. Sequenced cDNA libraries were mapped at the transcript and gene levels to the human genome (assembly GRCh38) using *Salmon* v0.8.2 (https://salmon.readthedocs.io/en/latest/salmon.html). We removed mapped genes for specific haemoglobin processes prior to data analysis.^26^ We normalised and compared gene expression using the R package DESeq2 was used to test for differential gene expression on log_2_ normalised gene counts.^27^ False Discovery Rate (FDR) corrected p-value <0.05 was used to threshold for significance in the differential gene expression analysis. All mapped genes were clustered per sample type using MIRU (https://kajeka.com) using correlation r^2^ >0.92 and MCL clustering tool,^28,29^ analysing clusters for biological enrichment using pathway over-representation analysis (ORA), in InnateDB (www.innatedb.com). Significantly enriched pathways were denoted by FDR corrected p-value <0.05. Network graphs of differentially expressed genes were generated using R package XGR (http://galahad.well.ox.ac.uk:3020/subneter/genes) and Gephi (https://gephi.org/), analysing graphs by network centrality.

Validated functional transcriptional gene response-modules were applied to CSF and blood data, module scores were calculated by geometric means of log_2_ normalised gene expression data per transcriptomic data set prior to all module analyses.^30,31^ Module scores were compared by outcome and sample type using Mann-Whitney U tests. Gene Set Enrichment Analysis (GSEA) was used to rank differentially expressed genes against the molecular signatures database for gene-ontology terms to analyse function-specific gene expression for individual cell types (http://software.broadinstitute.org/gsea/msigdb/index.jsp).

### Data Sharing

Mapped, sequence files for all included patients are available on a consent-basis through the European Phenome-Genome Archive at the European Bioinformatics Institute (EBI) https://www.ebi.ac.uk/ega/studies/EGAS00001003355

### Ethics

All participants or nominated guardians gave written informed consent for inclusion. Ethical approval for the transcriptomics study was granted by both the College of Medicine Research and Ethics Committee (COMREC), University of Malawi, (P.01/10/980, January 2011), and the Liverpool School of Tropical Medicine Research Ethics Committee, UK (P10.70, November 2010) Committee, Liverpool, UK.

## Findings

We extracted RNA of sequencing quality (*RIN* >7) from the blood of twenty-eight adults and paired CSF from thirteen adults with proven PM (Figure 1). The median age of the patients was 33 (range 26-66) years, mortality was 15/28 (52%) (Table 1), and 21/25 (84%) were HIV co-infected. All patients received parenteral ceftriaxone within 3 hours of arrival in hospital.^2^ Non-survivors of bacterial meningitis had lower Glasgow Coma Scores on admission to hospital OR 0·13 (95%CI 0·22 : 0·8, p=0·02). CSF white cell counts (CSF WCC) were equally low in both groups, but CSF bacterial loads were higher in non-survivors (p=0·02) (Table 1). Times from reported symptom onset to presentation did not differ between outcome groups.

**Figure 1:**
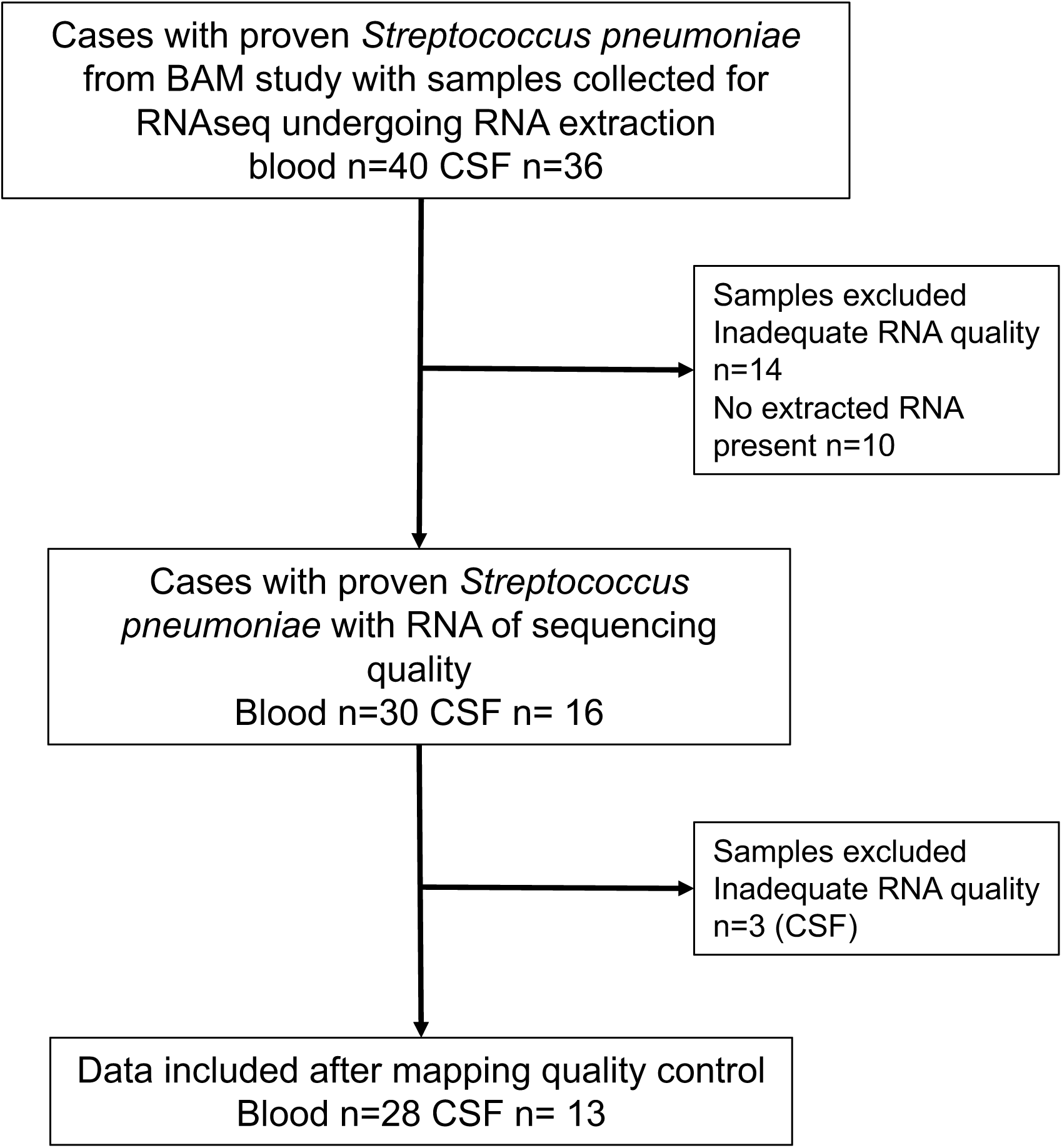
Selection of study patients for inclusion. BAM = Bundles for Adult Meningitis, CSF = cerebrospinal fluid RNA = Ribonucleic Acid.

**Table 1:** Demographic details of included patients

|  | Survivors<br>n=15 | Non-Survivors<br>n=13 | Univariate odds ratio<br>(95%CI)<br>*p*-value |
| --- | --- | --- | --- |
| **Median age in years (IQR)** | 32 (26-44) | 36 (26-46) | 1·0 (0·96 : 1·07)<br>0·75 |
| **Male Gender (n, %)** | 7 (46%) | 9 (69%) | 2·5 (0·54 : 12·1)<br>0·23 |
| **HIV co-infection (n, %)** | 12 (86%) | 10 (76%) | 0·25 (0·02 – 2·8)<br>0·26 |
| **Estimated BMI** | 22·4 (20·7-23·2) | 22·4 (19-23·5) | 0·90 (0·68 : 1·1)<br>0·44 |
| **Median CD4 count (cells/mm<sup>3</sup>) (IQR)** | 128 (25-290) | 255 (n=1) | 1·0 (0·98 : 1·04) |
| **Median Glasgow Coma Score (IQR)** | 14 (13-15) | 11 (6-13) | 0·13 (0·22 : 0·8)<br>0·02 |
| **Median CSF WCC (cells/mm<sup>3</sup>) (IQR)** | 28 (0-685) | 33 (7-209)<br>n=8 | 0·95 (0·4 : 2·2) 0·91 |
| **Median CSF bacterial load (DNA copies/ml) (IQR)** | 5·7E+05 (3206 – 4·7E+06) | 4·37E+07 (8·43 E+06 – 2·18E+08) | 6·0 (1·2 : 29·2) 0·02 |
| **Median Duration of symptoms at presentation in hrs (IQR)** | 48 (24-72) | 48 (21 – 60) | 0·99 (0·68 : 1·02) *p*=<br>0·66 |

We mapped all sequenced transcripts from both blood and CSF compartments at a gene level using *Salmon* and quantified transcripts using DESeq2. Global gene expression in the two compartments was examined using principal component analysis (PCA). Samples from the CSF transcriptome clustered separately to those from the blood transcriptome (Figure 2A), indicating important differences in gene expression between the two compartments. To investigate these differences, we ran all mapped genes expressed over 3 gene copies/million reads (CPM) in CSF and blood compartments through the clustering programme MIRU. Enriched gene clusters were tested against curated functional pathways compartments using InnateDB pathways over-representation analysis (ORA). The CSF compartment contained multiple gene clusters that were highly enriched for both innate and adaptive immune response pathways, damage repair and stress response, endothelial activation and synaptic neurotransmitter activity (Table 2). In contrast, the blood compartment contained fewer enriched gene clusters that mapped to immune pathways, and relatively greater enrichment of cell cycle and protein synthesis genes as well as gene clusters for hormonal signalling and carbohydrate metabolism (Table 3). Gene clusters were also detected in blood that mapped to endothelial damage, extracellular membrane breakdown and proteoglycan synthesis (Table 3).

**Figure 2:**
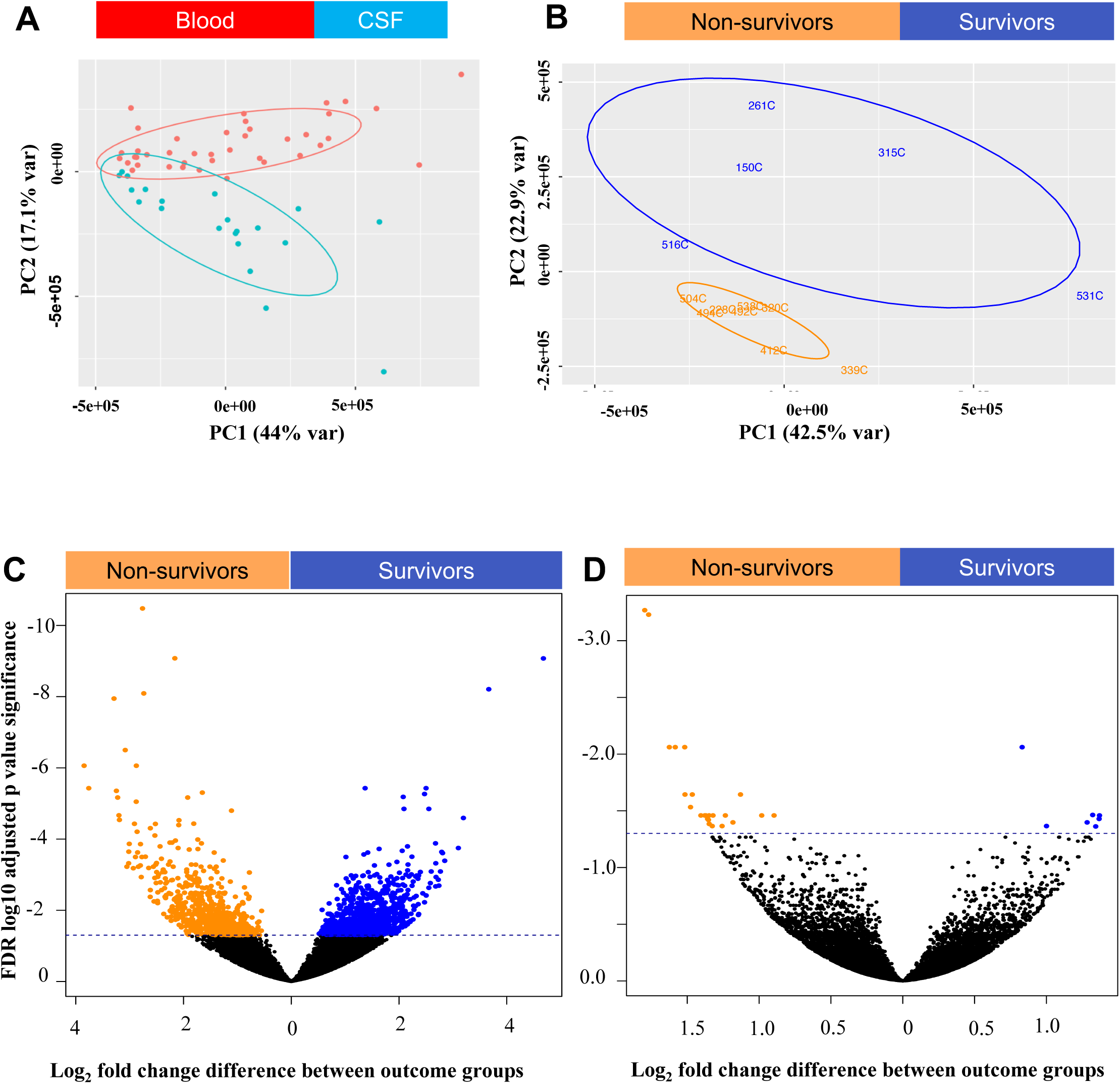
Transcriptional profiling with RNAseq reveals differential gene expression between survivors and non-survivors of pneumococcal meningitis, exclusively in the CSF compartment. Principal component analysis of gene expression in CSF (blue) and blood (red) compartments (**A**) and within in the CSF compartment between survivors (blue) and non-survivors (red) (**B**) of adults with proven pneumococcal meningitis (PM). Numbered dots represent individual patients, circles describe clusters. Depicted axes principal component 1 and principal component 2 account for the greatest variance between the groups. Volcano plot describing differential gene expression in CSF compartment (**C, n=13**) and blood (**D, n=28**) between survivors (upper right quadrant, blue)) and non-survivors (upper left quadrant, orange) in PM. Orange dots represent individual genes expressed over 1 log_2_ fold change differential expression (X axis) with adjusted log_10_ False Discovery Rate (FDR) p value (Y axis) <0.05 (horizontal dashed line).

**Table 2:**
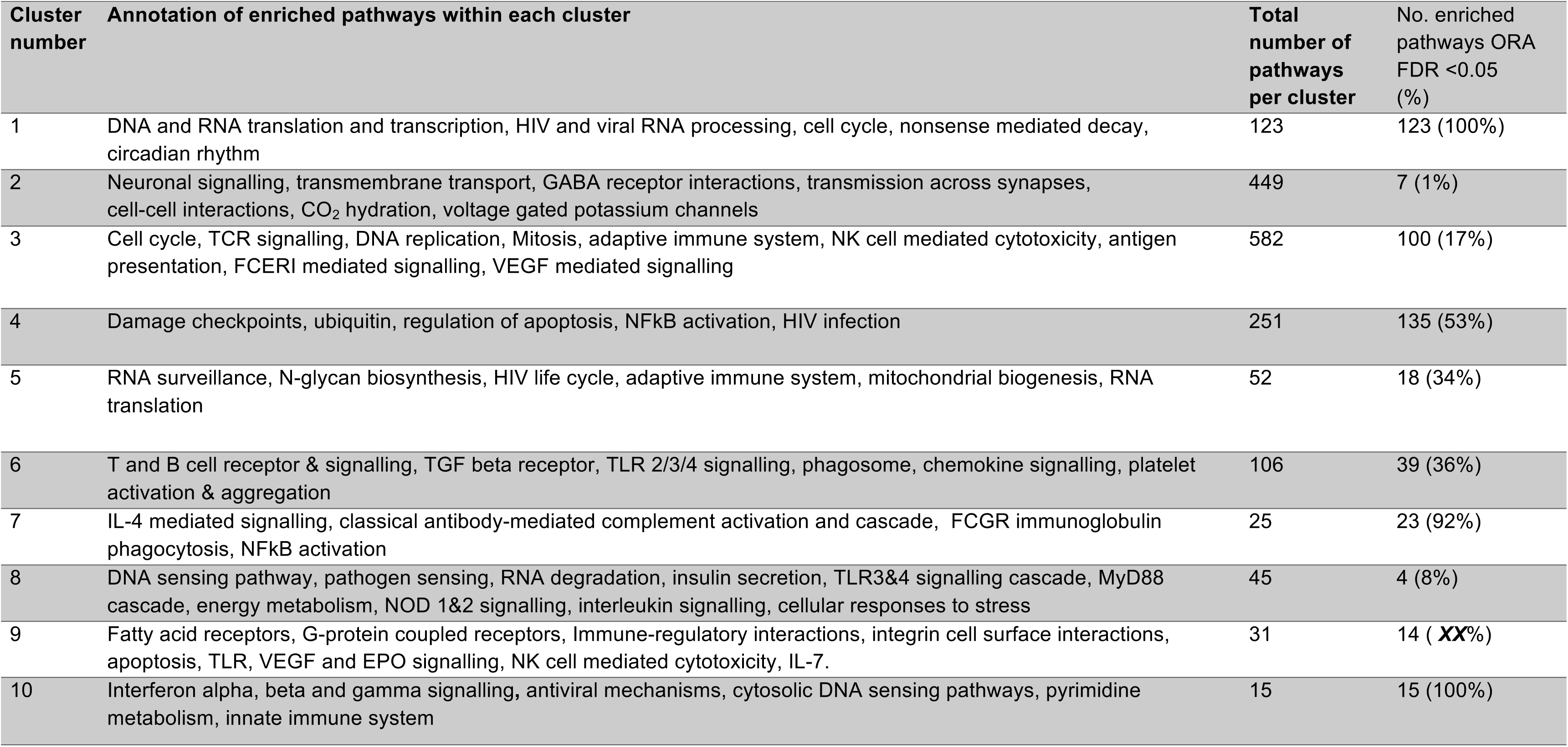
Summary of highly expressed gene clusters expressed in the CSF of adults with pneumococcal meningitis.

**Table 3:** Summary of highly expressed gene clusters expressed in the blood of adults with pneumococcal meningitis.

| Cluster number | Annotation of significantly enriched pathways within each cluster | Total number of pathways per cluster | No. enriched pathways ORA p-adj <0.05 (%) |
| --- | --- | --- | --- |
| 1 | Antigen processing: ubiquitination and proteasome degradation, haemoglobin chaperone | 88 | 8 (9%) |
| 2 | Antibody mediated complement activation, complement cascade, FCER1 activation of NF-κB, scavenging receptors, phagocytosis, B cell receptor signalling, scavenging heme from plasma | 23 | 23 (100%) |
| 3 | GABA receptor activation, synaptic transmission, axonal signalling, neuroactive ligand receptor activation, ECM proteoglycans, orphan transporters, membrane transport, degradation of ECM | 16 | 14 (87%) |
| 4 | DNA transcription, RNA translation, protein synthesis | 36 | 33 (91%) |
| 5 | Mitosis, antigen presentation, transcriptional control via p53& tp53, platelet production, DNA transcription, translation and repair | 69 | 65 (94%) |
| 6 | Chromatin modification and organisation | 4 | 3 (75%) |
| 7 | IL6, IL5, chemokine and cytokine signalling, BCR, FCGR dependent phagocytosis, EGFR signalling, FCGR activation, mRNA regulation, insulin signalling, cell-cell communication, interferon signalling, haemostasis, prolactin signalling | 37 | 24 (64%) |
| 8 | Ribosomal RNA, Eukaryotic and viral mRNA translation and transcription, protein synthesis, nonsense mediated decay, protein metabolism | 27 | 25 (92%) |
| 9 | IFN gamma, thrombin, IL6 signalling, platelet activation, GPCR signalling, TGF beta, membrane signalling, ubiquitin mediated proteolysis, TSH pathway, JAK/STAT pathway, regulation of actin cytoskeleton | 26 | 26 (16%) |
| 10 | Carbohydrate metabolism, TSH pathway | 6 | 6 (4%) |

We then tested if gene expression within each compartment differed between outcome groups. Principal component analysis of CSF gene expression showed a complete separation of sample clusters between survivors and non-survivors (Figure 2B). 1678 genes were differentially expressed (FDR <0.05 Log Fold Change (LFC) >0.5) in the CSF compartment; (762 genes upregulated in non-survivor CSF and 916 genes in survivor CSF) (Figure 1C). In contrast, very few genes were differentially expressed between outcome groups in the blood compartment (7 genes upregulated in survivor blood and 18 in non-survivor blood) (Figure 1D). We undertook a network analysis using a gene networking software (XGR) to determine if the differentially expressed genes were interconnected. Differentially expressed genes in survivor CSF clustered around two central gene hubs. The first represents genes involved in DNA and cellular response to damage and transcriptional control, including hubs *CCND1* and *HIST2H4A*. The second cluster represents genes involved in immunological signalling, including *PIK3CD* (multiple immunological signalling), *DVL2* and *FZD2* (Wnt signalling) and *FCGR3A* (Fc receptor for IgG) (Figure 3A). A further small gene cluster represented amino-acid metabolism. In comparison, genes enriched in the CSF from non-survivors clustered in a large central network of genes encoding pro-inflammatory mediators including *IL-1B*, *TNF*, *IL-6*, *MMP9* and *IGF-2* (Figure 3B). Peripheral clusters to this central network included genes involved in platelet aggregation, vasoconstriction and collagen breakdown, prostaglandin biosynthesis, cytoskeleton remodelling, and cell membrane synthesis (Figure 3B).

**Figure 3:**
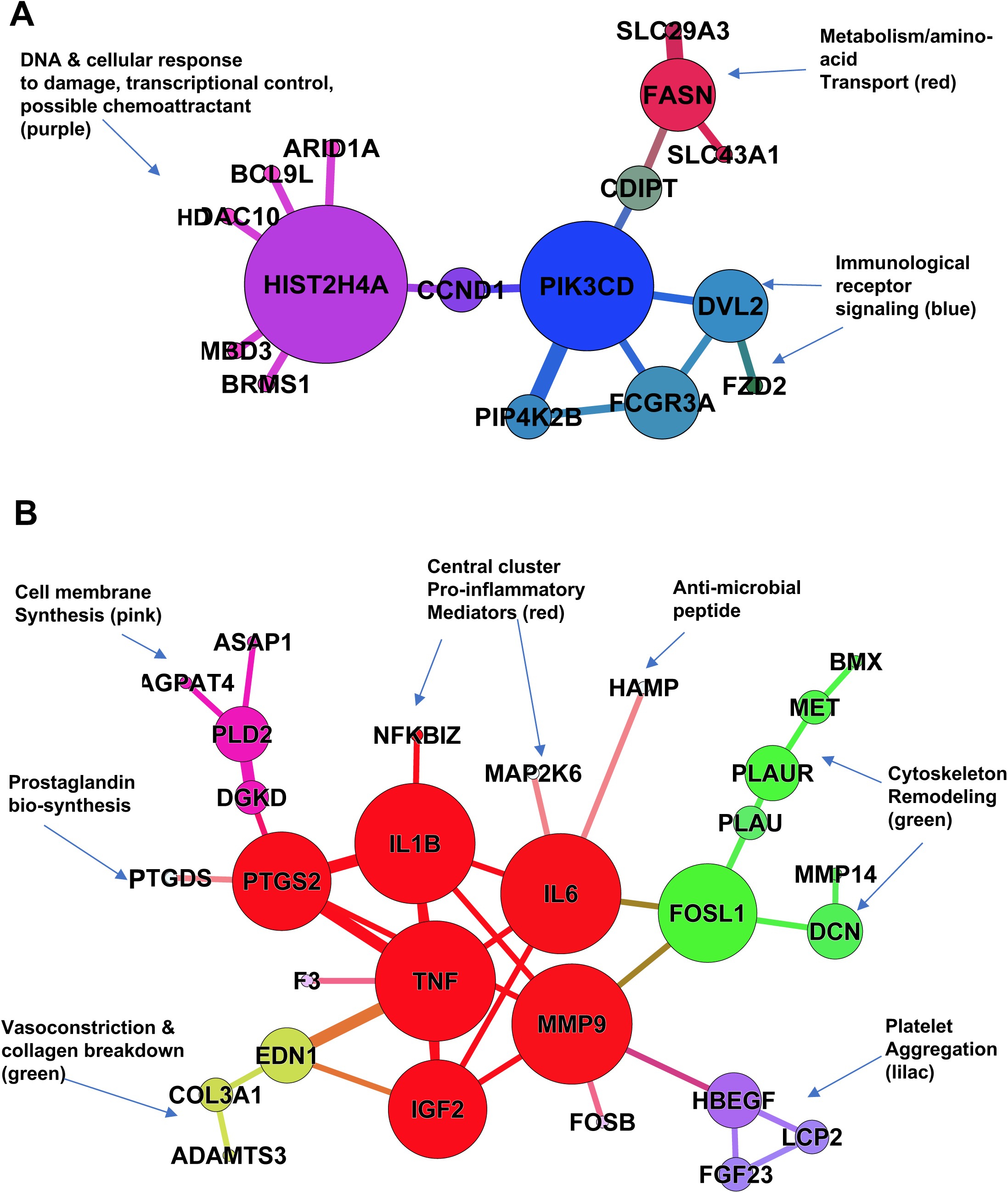
Comparison of the gene expression network analyses of differentially expressed genes reveals upregulation of transcriptional control in survivors and intense pro-inflammatory responses in non-survivors. Network analyses of significantly differentially expressed genes (FDR p-adj <0.05) in the CSF of survivors (A) and non-survivors (B) with pneumococcal meningitis. Gene clustering generated in XGR, graphs synthesised with Gephi, analysed with Wifan-Hu network centrality. Each node represents an individual gene, node size represents network centrality and connectivity. Edge thickness represents strength of the relationship between nodes. Colours represent individual clusters by function and connectivity.

To test the veracity of our findings, we re-analysed the mapped CSF data using an alternative software programme Gene Set Enrichment Analysis (GSEA). We used this software to annotate all differentially expressed genes for both outcome groups with Gene Ontology (GO) functional terms. GO terms that were highly enriched in the GSEA annotation (FDR <0.05) included ‘Positive regulation of reactive oxygen species’, ‘Extracellular matrix’ and ‘Granulocyte migration’ in non-survivors. ‘DNA replication’, ‘Damaged DNA binding’ and ‘G-protein coupled chemoattractant’ were enriched in survivors, amongst others (Supplementary data 1).

To determine the transcriptional foot-print of both cell-specific gene expression and functional inflammatory processes, we compared the expression of transcriptional response-modules for specific white cell subsets and cytokines between survivors and non-survivors in CSF and blood, validated in patients in sub-Saharan Africa.^30,31^ The geometric mean expression of the neutrophil, monocyte, T-cell, TNF-and Type 2 interferon-response modules did not differ between survivors and non-survivors in the CSF transcriptome (Figure 4). However, expression of IL-17, IL-10 and Type 1 Interferon gene response modules was significantly increased in the CSF of non-survivors compared to survivors (Figure 4). No differences in cell or functional gene response-module expression between outcome groups were detected in the blood transcriptome, apart from increased expression of the monocyte response-module in survivors (Supplementary Figure 1).

**Figure 4:**
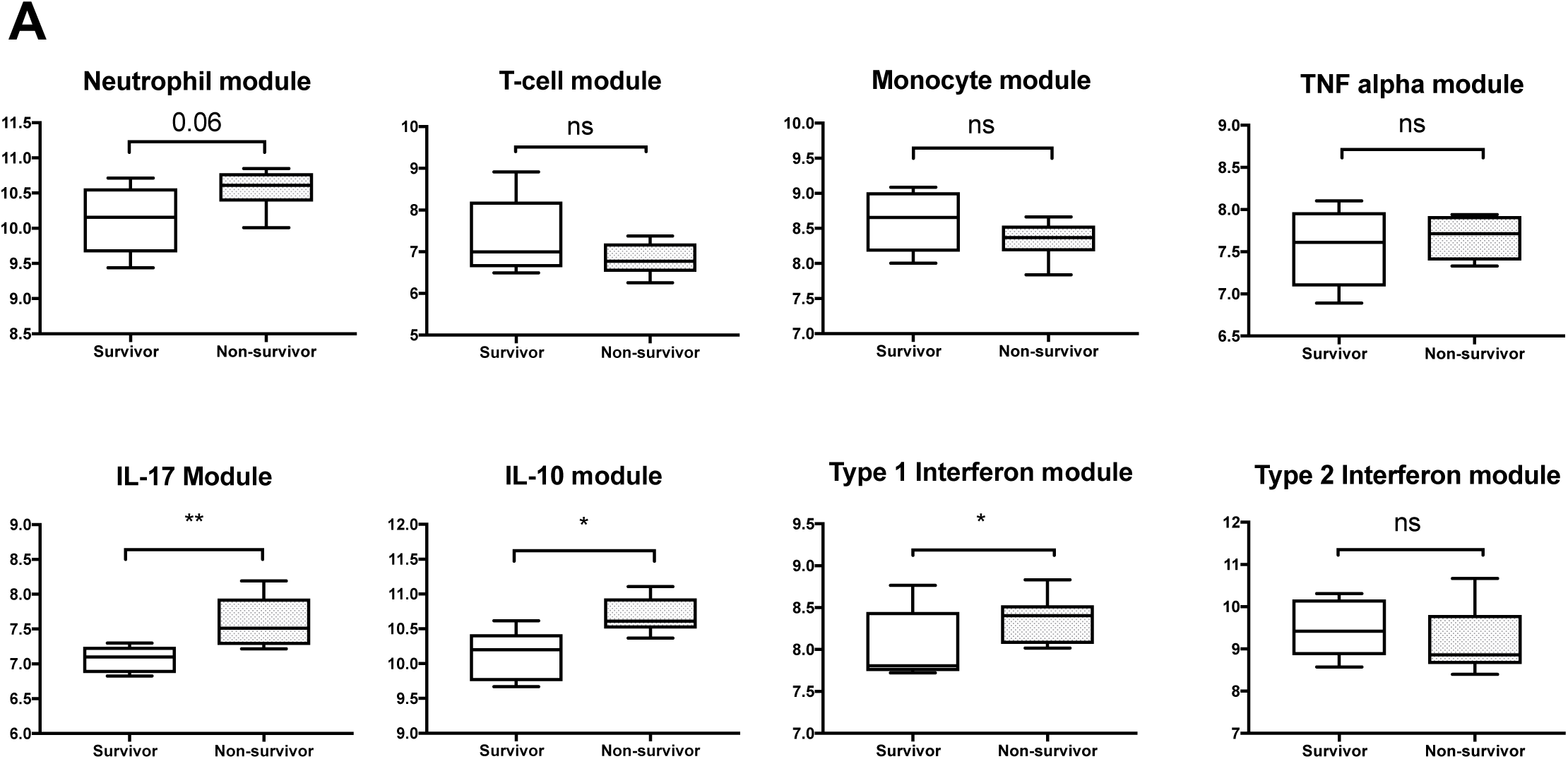
The inflammatory response in non-survivors of pneumococcal meningitis is associated predominately with upregulation of functional of pro-inflammatory cytokine gene response modules, but not cell-specific modules within the CSF compartment. Individual CSF module scores per patient by outcome group. Module scores (y axis) were calculated by geometric mean of log_2_ transformed gene transcript counts per million (CPM) reads across all genes in each module. Boxes show geometric mean with 95% confidence intervals, whiskers indicate range. Clinical outcome at 40 days post presentation with proven meningitis. Statistical significance calculated by Mann-Whitney U test, p<0.05 determined statistical significance*=p<0.05, **=p<0.01, *** p<0.001

Neutrophils were the predominant cell type (up to 95%) in the CSF of patients with PM. Low CSF leukocyte count is strongly associated with outcome, in this study CSF leukocyte counts were not statistically different between the outcome groups^2,19^ The neutrophil gene response-module correlates with absolute neutrophil count and activity, and as such reflects the CSF leukocyte counts in our study.^30^ However, the response-module does not test the activity of genes for individual functions such as activation, chemotaxis or phagocytosis. We hypothesised that amongst the CSF neutrophil population, cells may exist in different functional states. To examine expression of specific neutrophil functions, we used GSEA leading edge analysis to test for enrichment of Gene Ontology (GO) functional neutrophil terms in the CSF transcriptome between outcome groups (Supplementary data 1, Supplementary Figure 2). To consolidate these findings, we then used the modular analytical approach to test the expression of genes from individual neutrophil-specific functional GO terms. We found enrichment of genes associated with neutrophil chemotaxis and apoptosis in non-survivor CSF, but no differences in expression of genes for either neutrophil degranulation or cytotoxicity (Figure 5A, Supplementary data 2). Neutrophils expressing persistence or survival-associated genes *CSF3* and *NR4A3/1* have been associated with pro-inflammatory states in models of neutrophilic inflammation.^32,33^ To test for evidence of neutrophil survival/persistence, we examined individual expression of these genes. Expression of *CSF3*, *NR4A3* and *NR4A1* were all markedly enriched in non-survivor CSF (Figure 5B), suggesting the presence of a sub-population of neutrophils within the CSF expressing a strongly pro-inflammatory phenotype.

**Figure 5:**
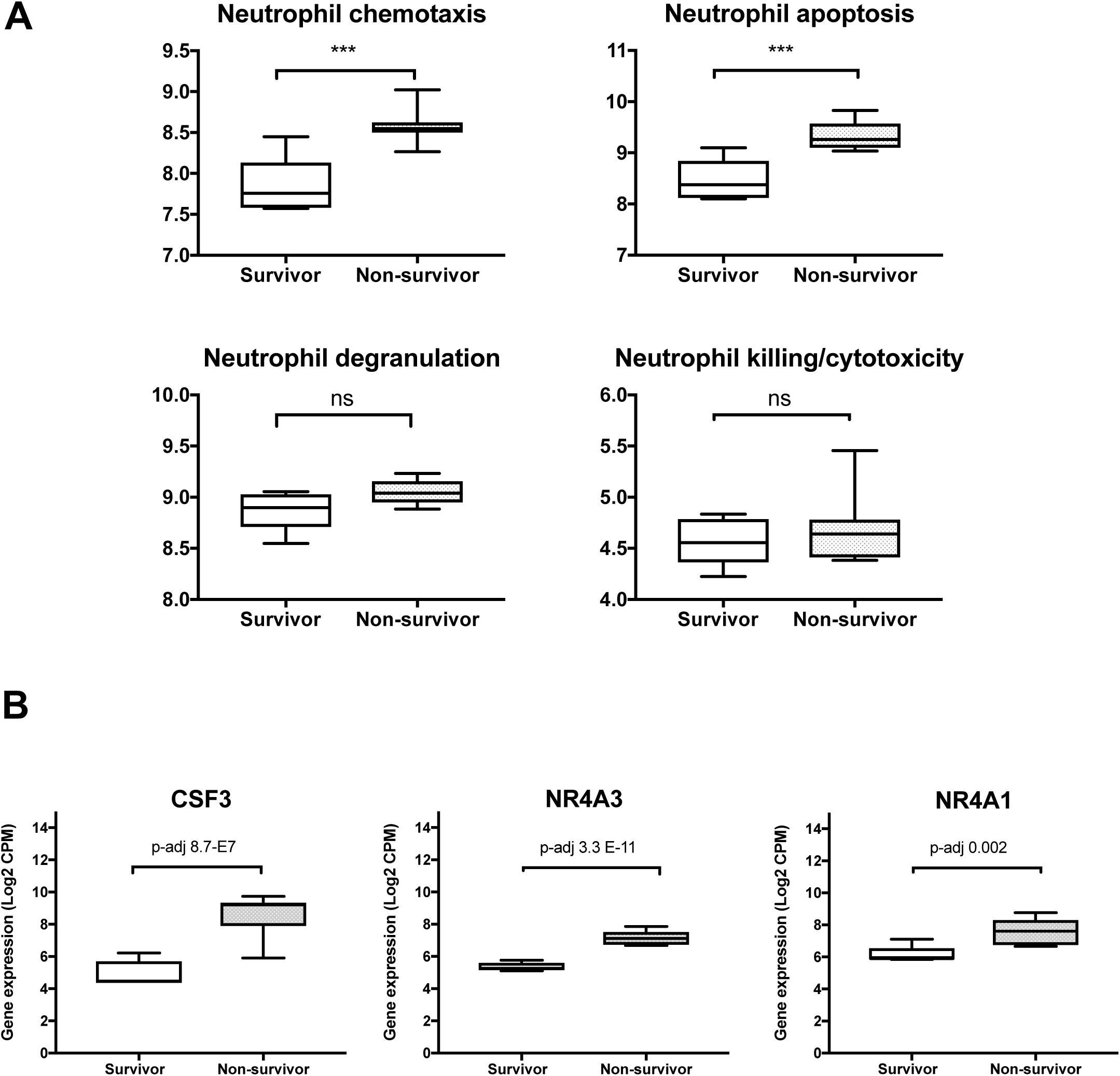
Expression of neutrophil associated transcripts in the CSF of adults with pneumococcal meningitis shows upregulation of genes for neutrophil chemotaxis, apoptosis and persistence/survival, but not active killing in non-survivors. Expression of neutrophil related Gene Ontology (GO) terms by outcome group (A) and expression of individual neutrophil specific persistence/survival genes (B). CSF3 = Colony Stimulating Factor 3, NR4A3 and NR4A1 = Nuclear Receptor subfamily 4 group A member 3/1. Log_2_ gene counts per million reads (CPM) per patient calculated across all genes in GO term is shown on the Y axis. Boxes show geometric mean with 95% confidence intervals, whiskers indicate range. Differences between outcome groups calculated using univariate non-parametric testing. p<0.05 determined statistical significance. Results validated using Gene Set Expression Analysis (GSEA), data in supplementary.*=p<0.05, **=p<0.01, *** p<0.001

## Interpretation

Why patients with PM in LMICs with high HIV-1 prevalence have worse clinical outcomes compared to better resourced settings remains poorly understood. Our transcriptional approach has identified the harmful host inflammatory pathways that characterise non-surviving adults with PM in Malawi, going some way towards explaining why interventions targeted on attenuating the inflammatory response have failed. Patients enrolled in our study were typical of PM adults in sub-Saharan Africa: predominately young, HIV-1 co-infected, and presenting with profoundly low CSF neutrophil counts and very high bacterial loads.^1,34^ The differences in pro-inflammatory transcriptomic responses between survivors and non-survivors were solely in the CSF compartment. The inflammatory CSF cascade in non-survivors of PM was dominated by neutrophil dysfunction and a cluster of transcripts coding for pro-inflammatory mediators including IL-17 and Type 1 interferon genes. This contrasts with the increase in expression of genes linked to transcriptional regulation, damage repair and cell-cell signalling in survivors. Many of the upregulated gene transcripts in the CSF of non-survivors correspond to CSF proteins associated with poor outcome from meningitis identified in a number of different studies, including potent tissue collagenases, matrix-metalloproteinases MMP 8&9.^35-40^

Both IL-17 and T1-IFN driven mechanisms are important components of the mucosal host response to *S. pneumoniae*, and may be associated with neutrophil recruitment across the blood brain barrier. IL-17 activity in the CSF is a new finding in adults with PM;^46^; this cytokine has potent downstream activity including induction of tissue collagenases (e.g. MMP-8 and 9) and vaso-active mediators (e.g. VEGF) in infected tissue, facilitating neutrophil recruitment and enhancing phagocytosis in infected tissue.^39,47,48^ T1-IFN activity represents a group of common pro-inflammatory cytokines that are particularly important in the inflammatory response to both viral and bacterial infections, including HIV-1. Upregulation of IL-17 and T1-IFN module genes and expression of downstream mediators, including MMP-8 and 9, suggests that protective mucosal responses against *S.pneumoniae* may be damaging to brain cells.

The CSF bacterial load was higher in non-survivors compared to survivors in our study, presenting a potent stimulus for the upregulated T1-IFN and IL-17 driven pro-inflammatory cascade.^49,50^ We have previously shown in earlier, larger studies that persistence of pneumolysin in the CSF rather than absolute CSF bacterial load is more strongly associated with poor outcome in PM.^51,52^ Pneumolysin is a pneumococcal pore-forming toxin and possible TLR4 agonist, and is involved in IL-17 activation of the neutrophil NLRP3 inflammasome in models of pneumococcal infection.^53^ Pneumolysin has been shown in PM to kill neutrophils, attenuate CNS leukocyte counts and cause direct neuronal cell death through synaptic dysfunction.^14,54,55^ Our data show that the presence of increased pneumococci in the CSF is a significant driver of the observed CSF pro-inflammatory cascade, over-expression of the IL-17 pathway, attenuated neutrophil activity and increased host cell damage in may be due to correspondingly increased production of pneumolysin in the CSF.^56^

High bacterial loads in the CSF of non-survivors in our study suggest bacterial control by neutrophils may be ineffective in these patients, we hypothesised that the neutrophil transcriptome would be down-regulated in non-survivors. Neutrophils are both the most abundant CSF cell type in PM, and a critical component of the innate host response to infection in bacterial meningitis.^12^ Low CSF leukocyte counts are recognised as a poor prognostic factor in bacterial meningitis in all settings.^9,19,40^ Interestingly, we observed increased CSF neutrophil chemotaxis signalling in the presence of the pro-inflammatory cascade in non-survivors, but we did not detect the expected increases in either CSF neutrophil count, neutrophil response module expression or gene expression for degranulation or phagocytosis that should follow from expression of these powerful migration signals.^57,58^ However, we detected significant upregulation in non-survivors of genes coding for neutrophil persistence and survival including *CSF3/NR4A3/NR4A1*. Expression of these genes may be triggered by cytokine release from necrotic neutrophils,^33^ and are associated with more severe inflammation and tissue damage in other tissues^32,33,59^ including through increased production of IL-17 and TNF alpha from surrounding cells.^49,60^ The relatively reduced expression of neutrophil genes related to phagocytosis, neutrophil killing and degranulation in the context of increased neutrophil migration signalling in non-survivors of PM may reflect poor functional activity of these persisting neutrophils. Hence it is possible that neutrophil damage, mediated by pneumolysin, causes functional neutrophil failure leading to a pro-inflammatory burden of neutrophil persistence and necrosis in the CSF that underpins many of the associations between inflammatory gene expression and poor outcome in our study.

Adjunctive treatment with broadly effective anti-inflammatory agents such as dexamethasone have failed to improve the poor outcome of PM in LMICs.^16,61^ Dexamethasone is relatively ineffective against IL-17 mediated neutrophilic inflammation.^62,63^ Selective blockade of the IL-17 and T1-IFN pathways identified in this study may limit the damaging elements of the inflammatory cascade^32,64,65^ but given the complex inter-relationships that we have demonstrated, these may have unpredictable effects on outcome.^66,67^ The specific roles of these mediators in active disease needs to be elucidated before clinical trials of any of these agents can be undertaken.

### Limitations

There are several limitations to our study. Included patients were predominately HIV-1 co-infected, comparisons by HIV serostatus were not possible due to small numbers of HIV-negative patients. We were unable to stratify our analysis by viral load or CD4 count as these were not included in the original trial protocol. All HIV-1 co-infected patients were WHO clinical stage 3. The relatively small numbers of patients with high quality CSF RNAseq libraries limited our ability to stratify transcriptome signatures by either neutrophil count or bacterial load. Specific transcriptional modules for critical neutrophil functions, such as phagocytosis and trans-endothelial migration are currently lacking, instead we used the gene response-module analysis approach using GO terms that have high sensitivity but untested specificity. Sufficient CSF was not available to validate our findings at the protein level; but our findings are supported by other biomarker studies.^39,40,48^ Prehospital delay may have influenced the differences in transcriptional responses seen between survivors and non-survivors, although estimates of pre-hospital disease onset times were not associated with outcome in either our study or in a larger Malawi meningitis database.^2,10^

### Conclusions

Our comparative analysis of the CSF transcriptome between survivors and non-survivors of proven PM implicates both upregulation of protective mucosal responses and pro-inflammatory persistent neutrophil genes with damaging inflammation in the CNS of non-survivors. Understanding the triggers for neutrophil-driven inflammation will be critical to developing effective and targeted interventions for this devastating disease.

## Supporting information

## Declaration of Interests

All authors have no conflicts of interest to declare

## Funding

This study was funded by a Clinical Lecturer Starter Grant from the Academy of Medical Sciences (UK), by a PhD Fellowship in Global Health to EW from the Wellcome Trust (089671/B/09/Z) and by a Postdoctoral Clinical Research Fellowship to GP from the Wellcome Trust (WT101766). The Malawi-Liverpool-Wellcome Trust Clinical Research Programme is supported by a core grant from the Wellcome Trust (101113/Z/13/Z). This work was undertaken at UCLH/UCL who received a proportion of funding from the National Institute for Health Research University College London Hospitals Department of Health’s NIHR Biomedical Research Centre. JSB, MN and CV are supported by the Centre’s funding scheme. The funders of the study had no role in study design, data collection, data analysis, data interpretation, or writing of the report. The corresponding author had full access to all the data and the final responsibility to submit for publication.

## Acknowledgements

The authors would like to thank the study patients and guardians, the Bundles for Adult Meningitis (BAM) research team, clinical and laboratory staff at the Queen Elizabeth Central Hospital and Malawi-Liverpool-Wellcome Trust Clinical Research Programme in Blantyre Malawi for support given during the study and Professor Mike Levin and Dr Victoria Wright of Imperial College UK for support at the early stages of the project. We would like to thank Professor Judith Breuer and the staff of the Pathogen Genomics Unit at University College London for their assistance with library preparation and RNA sequencing. The authors acknowledge the use of the UCL Legion High Performance Computing Facility (Legion@UCL), and associated support services, in the completion of this work.

## Supplementary Data, Tables and Figures

**Supplementary Figure 1:**
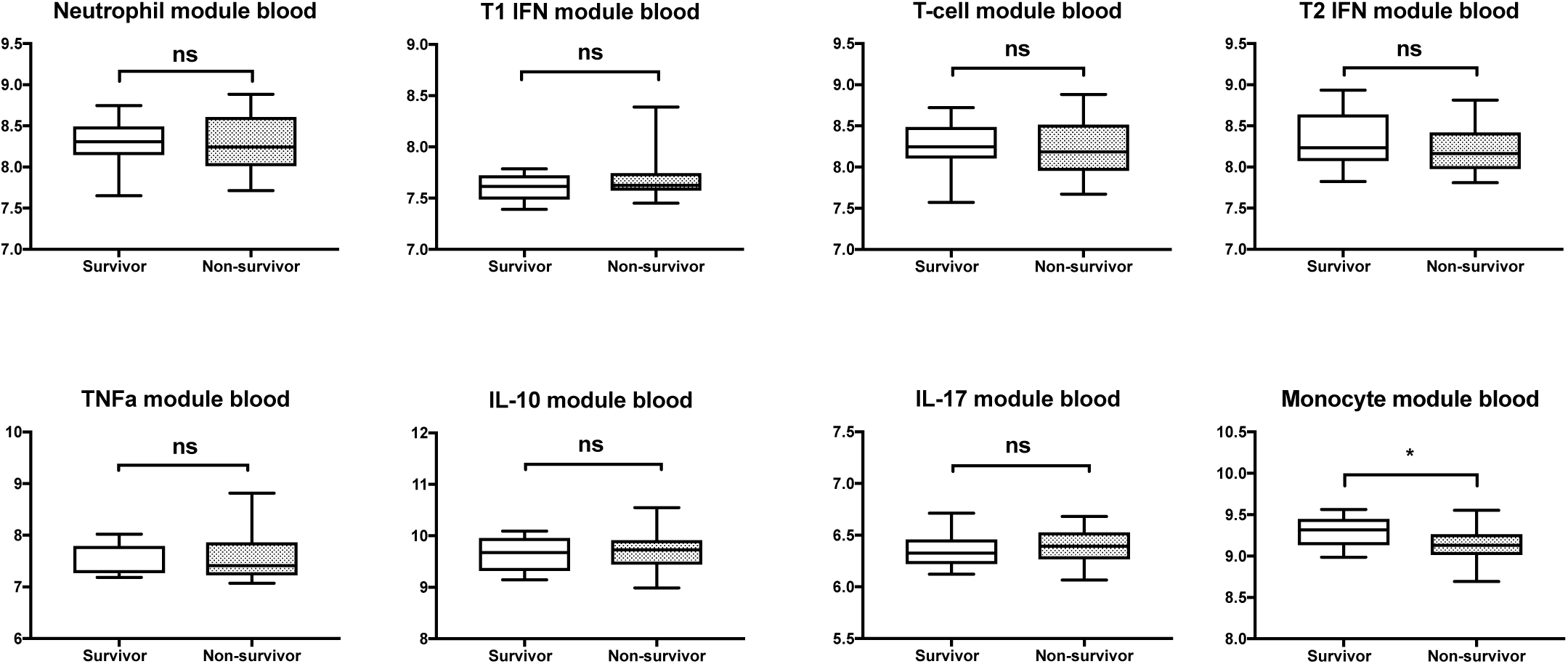
Transcriptional module expression in blood of patients with pneumococcal meningitis shows no difference in the transcriptional foot print in the blood compartment between survivors and non-survivors, with the exception of increased expression of monocyte activity in survivors. Individual blood gene expression module scores per patient by outcome group. Module scores (y axis) were calculated by geometric mean of log_2_ transformed gene transcript counts per million (CPM) reads across all genes in each module. Boxes show geometric mean with 95% confidence intervals, whiskers indicate range. Clinical outcome at 40 days post presentation with proven meningitis. Statistical significance calculated by Mann-Whitney U test, p<0.05 determined statistical significance*=p<0.05, **=p<0.01, *** p<0.001

**Supplementary Figure 2:**
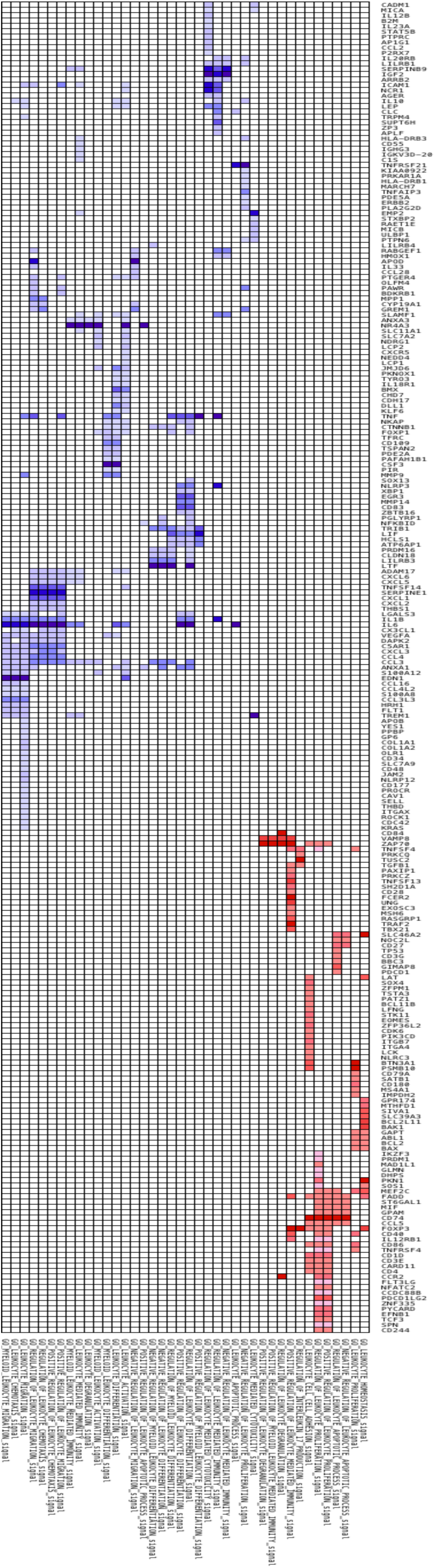
Enrichment of individual genes associated with leukocyte specific GO terms in the CSF of adults with PM demonstrates differences in neutrophil activity between outcome groups. Heatmap showing Enrichment of GO terms (FDR 0.05) on GSEA analysis sub-setted for leukocyte specific enrichment are shown on the X axis. Individual genes included within each GO term are shown on the y axis. Genes enriched in non-survivor CSF are shown in red, survivor CSF in blue. Colour intensity relates to differential expression between the two outcome groups.

**Supplementary Data 1**: CSF mapped gene expression data

**Supplementary Data 2**: GSEA output for CSF transcriptional data mapped against

## References

1. Group GBDNDC. Global, regional, and national burden of neurological disorders during 1990-2015: a systematic analysis for the Global Burden of Disease Study 2015. Lancet neurology 2017.

2. Wall EC, Mukaka M, Denis B, et al. Goal directed therapy for suspected acute bacterial meningitis in adults and adolescents in sub-Saharan Africa. PloS one 2017; 12(10): e0186687.

3. Collaborators GBDCoD. Global, regional, and national age-sex specific mortality for 264 causes of death, 1980-2016: a systematic analysis for the Global Burden of Disease Study 2016. Lancet 2017; 390(10100): 1151–210.

4. Gessner BD, Mueller JE, Yaro S. African meningitis belt pneumococcal disease epidemiology indicates a need for an effective serotype 1 containing vaccine, including for older children and adults. BMC infectious diseases 2010; 10: 22.

5. Kwarteng A, Amuasi J, Annan A, et al. Current meningitis outbreak in Ghana: Historical perspectives and the importance of diagnostics. Acta Trop 2017; 169: 51–6.

6. Wall EC, Everett DB, Mukaka M, et al. Bacterial meningitis in Malawian adults, adolescents, and children during the era of antiretroviral scale-up and Haemophilus influenzae type b vaccination, 2000-2012. Clin Infect Dis 2014; 58(10): e137–45.

7. Okike IO, Ribeiro S, Ramsay ME, Heath PT, Sharland M, Ladhani SN. Trends in bacterial, mycobacterial, and fungal meningitis in England and Wales 2004-11: an observational study. The Lancet infectious diseases 2014; 14(4): 301–7.

8. Bijlsma MW, Brouwer MC, Kasanmoentalib ES, et al. Community-acquired bacterial meningitis in adults in the Netherlands, 2006-14: a prospective cohort study. The Lancet infectious diseases 2016; 16(3): 339–47.

9. van de Beek D, de Gans J, Spanjaard L, Weisfelt M, Reitsma JB, Vermeulen M. Clinical features and prognostic factors in adults with bacterial meningitis. The New England journal of medicine 2004; 351(18): 1849–59.

10. Wall EC, Cartwright K, Scarborough M, et al. High Mortality amongst Adolescents and Adults with Bacterial Meningitis in Sub-Saharan Africa: An Analysis of 715 Cases from Malawi. PloS one 2013; 8(7): e69783.

11. Wang S, Peng L, Gai Z, et al. Pathogenic Triad in Bacterial Meningitis: Pathogen Invasion, NF-kappaB Activation, and Leukocyte Transmigration that Occur at the Blood-Brain Barrier. Frontiers in microbiology 2016; 7: 148.

12. Mook-Kanamori BB, Geldhoff M, van der Poll T, van de Beek D. Pathogenesis and pathophysiology of pneumococcal meningitis. Clinical microbiology reviews 2011; 24(3): 557–91.

13. Doran KS, Fulde M, Gratz N, et al. Host-pathogen interactions in bacterial meningitis. Acta neuropathologica 2016; 131(2): 185–209.

14. Wippel C, Maurer J, Fortsch C, et al. Bacterial Cytolysin during Meningitis Disrupts the Regulation of Glutamate in the Brain, Leading to Synaptic Damage. PLoS pathogens 2013; 9(6): e1003380.

15. de Gans J, van de Beek D. Dexamethasone in adults with bacterial meningitis. The New England journal of medicine 2002; 347(20): 1549–56.

16. Scarborough M, Gordon SB, Whitty CJ, et al. Corticosteroids for bacterial meningitis in adults in sub-Saharan Africa. N Engl J Med 2007; 357(24): 2441–50.

17. Ajdukiewicz KM, Cartwright KE, Scarborough M, et al. Glycerol adjuvant therapy in adults with bacterial meningitis in a high HIV seroprevalence setting in Malawi: a double-blind, randomised controlled trial. The Lancet infectious diseases 2011; 11(4): 293–300.

18. Schut ES, Brouwer MC, Scarborough M, et al. Validation of a Dutch risk score predicting poor outcome in adults with bacterial meningitis in Vietnam and Malawi. PloS one 2012; 7(3): e34311.

19. Wall EC, Mukaka M, Scarborough M, et al. Prediction of Outcome From Adult Bacterial Meningitis in a High-HIV-Seroprevalence, Resource-Poor Setting Using the Malawi Adult Meningitis Score (MAMS). Clinical infectious diseases : an official publication of the Infectious Diseases Society of America 2017; 64(4): 413–9.

20. van Veen KE, Brouwer MC, van der Ende A, van de Beek D. Bacterial meningitis in patients with HIV: A population-based prospective study. The Journal of infection 2016; 72(3): 362–8.

21. Davenport EE, Burnham KL, Radhakrishnan J, et al. Genomic landscape of the individual host response and outcomes in sepsis: a prospective cohort study. Lancet Respir Med 2016; 4(4): 259–71.

22. Burnham KL, Davenport EE, Radhakrishnan J, et al. Shared and Distinct Aspects of the Sepsis Transcriptomic Response to Fecal Peritonitis and Pneumonia. American journal of respiratory and critical care medicine 2017; 196(3): 328–39.

23. Dunning J, Blankley S, Hoang LT, et al. Progression of whole-blood transcriptional signatures from interferon-induced to neutrophil-associated patterns in severe influenza. Nature immunology 2018; 19(6): 625–35.

24. Banerjee A, Van Sorge NM, Sheen TR, Uchiyama S, Mitchell TJ, Doran KS. Activation of brain endothelium by pneumococcal neuraminidase NanA promotes bacterial internalization. Cellular microbiology 2010; 12(11): 1576–88.

25. Mahdi LK, Wang H, Van der Hoek MB, Paton JC, Ogunniyi AD. Identification of a novel pneumococcal vaccine antigen preferentially expressed during meningitis in mice. The Journal of clinical investigation 2012; 122(6): 2208–20.

26. Westermann AJ, Gorski SA, Vogel J. Dual RNA-seq of pathogen and host. Nature reviews Microbiology 2012; 10(9): 618–30.

27. Anders S, Huber W. Differential expression analysis for sequence count data. Genome biology 2010; 11(10): R106.

28. Theocharidis A, van Dongen S, Enright AJ, Freeman TC. Network visualization and analysis of gene expression data using BioLayout Express(3D). Nat Protoc 2009; 4(10): 1535–50.

29. van Dongen S. A cluster algorithm for graphs. Amsterdam, The Netherlands: National Research Institute for Mathematics and Computer Sciences, 2000.

30. Bell LC, Pollara G, Pascoe M, et al. In Vivo Molecular Dissection of the Effects of HIV-1 in Active Tuberculosis. PLoS pathogens 2016; 12(3): e1005469.

31. Pollara G, Murray MJ, Heather JM, et al. Validation of Immune Cell Modules in Multicellular Transcriptomic Data. PloS one 2017; 12(1): e0169271.

32. Prince LR, Prosseda SD, Higgins K, et al. NR4A orphan nuclear receptor family members, NR4A2 and NR4A3, regulate neutrophil number and survival. Blood 2017; 130(8): 1014–25.

33. Thompson AA, Elks PM, Marriott HM, et al. Hypoxia-inducible factor 2alpha regulates key neutrophil functions in humans, mice, and zebrafish. Blood 2014; 123(3): 366–76.

34. Edmond K, Clark A, Korczak VS, Sanderson C, Griffiths UK, Rudan I. Global and regional risk of disabling sequelae from bacterial meningitis: a systematic review and meta-analysis. The Lancet infectious diseases 2010; 10(5): 317–28.

35. Roine I, Pelkonen T, Bernardino L, et al. Predictive value of cerebrospinal fluid matrix metalloproteinase-9 and tissue inhibitor of metalloproteinase-1 concentrations in childhood bacterial meningitis. The Pediatric infectious disease journal 2014; 33(7): 675–9.

36. Tsai H-C, Liu S-F, Wu K-S, et al. Dynamic changes of matrix metalloproteinase-9 in patients with Klebsiella pneumoniae meningitis. Inflammation 2008; 31(4): 247–53.

37. Barichello T, Generoso JS, Simoes LR, et al. Interleukin-1beta Receptor Antagonism Prevents Cognitive Impairment Following Experimental Bacterial Meningitis. Curr Neurovasc Res 2015; 12(3): 253–61.

38. Grandgirard D, Gaumann R, Coulibaly B, et al. The causative pathogen determines the inflammatory profile in cerebrospinal fluid and outcome in patients with bacterial meningitis. Mediators of inflammation 2013; 2013: 312476.

39. Mankhambo LA, Banda DL, Group IPDS, et al. The role of angiogenic factors in predicting clinical outcome in severe bacterial infection in Malawian children. Crit Care 2010; 14(3): R91.

40. Geldhoff M, Mook-Kanamori BB, Brouwer MC, et al. Inflammasome activation mediates inflammation and outcome in humans and mice with pneumococcal meningitis. BMC infectious diseases 2013; 13: 358.

41. Freundt-Revilla J, Maiolini A, Carlson R, et al. Th17-skewed immune response and cluster of differentiation 40 ligand expression in canine steroid-responsive meningitis-arteritis, a large animal model for neutrophilic meningitis. Journal of neuroinflammation 2017; 14(1): 20.

42. Pido-Lopez J, Kwok WW, Mitchell TJ, Heyderman RS, Williams NA. Acquisition of pneumococci specific effector and regulatory Cd4+ T cells localising within human upper respiratory-tract mucosal lymphoid tissue. PLoS pathogens 2011; 7(12): e1002396.

43. Peno C, Banda DH, Jambo N, et al. Alveolar T-helper 17 responses to streptococcus pneumoniae are preserved in ART-untreated and treated HIV-infected Malawian adults. The Journal of infection 2018; 76(2): 168–76.

44. Campos IB, Herd M, Moffitt KL, et al. IL-17A and complement contribute to killing of pneumococci following immunization with a pneumococcal whole cell vaccine. Vaccine 2017; 35(9): 1306–15.

45. Wright AK, Bangert M, Gritzfeld JF, et al. Experimental human pneumococcal carriage augments IL-17A-dependent T-cell defence of the lung. PLoS pathogens 2013; 9(3): e1003274.

46. Morichi S, Urabe T, Morishita N, et al. Pathological analysis of children with childhood central nervous system infection based on changes in chemokines and interleukin-17 family cytokines in cerebrospinal fluid. J Clin Lab Anal 2018; 32(1).

47. Veldhoen M. Interleukin 17 is a chief orchestrator of immunity. Nature immunology 2017; 18(6): 612–21.

48. Roine I, Pelkonen T, Lauhio A, et al. Changes in MMP-9 and TIMP-1 Concentrations in Cerebrospinal Fluid after 1 Week of Treatment of Childhood Bacterial Meningitis. Journal of clinical microbiology 2015; 53(7): 2340–2.

49. Ritchie ND, Ritchie R, Bayes HK, Mitchell TJ, Evans TJ. IL-17 can be protective or deleterious in murine pneumococcal pneumonia. PLoS pathogens 2018; 14(5): e1007099.

50. Damjanovic D, Khera A, Medina MF, et al. Type 1 interferon gene transfer enhances host defense against pulmonary Streptococcus pneumoniae infection via activating innate leukocytes. Mol Ther Methods Clin Dev 2014; 1: 5.

51. Wall EC, Gordon SB, Hussain S, et al. Persistence of pneumolysin in the cerebrospinal fluid of patients with pneumococcal meningitis is associated with mortality. Clinical infectious diseases : an official publication of the Infectious Diseases Society of America 2012; 54(5): 701–5.

52. Wall EC, Gritzfeld JF, Scarborough M, et al. Genomic pneumococcal load and CSF cytokines are not related to outcome in Malawian adults with meningitis. The Journal of infection 2014; 69(5): 440–6.

53. Hassane M, Demon D, Soulard D, et al. Neutrophilic NLRP3 inflammasome-dependent IL-1beta secretion regulates the gammadeltaT17 cell response in respiratory bacterial infections. Mucosal Immunol 2017; 10(4): 1056–68.

54. Jim KK, Engelen-Lee J, van der Sar AM, et al. Infection of zebrafish embryos with live fluorescent Streptococcus pneumoniae as a real-time pneumococcal meningitis model. Journal of neuroinflammation 2016; 13(1): 188.

55. Ullah I, Ritchie ND, Evans TJ. The interrelationship between phagocytosis, autophagy and formation of neutrophil extracellular traps following infection of human neutrophils by Streptococcus pneumoniae. Innate Immun 2017; 23(5): 413–23.

56. Penaloza HF, Nieto PA, Munoz-Durango N, et al. Interleukin-10 plays a key role in the modulation of neutrophils recruitment and lung inflammation during infection by Streptococcus pneumoniae. Immunology 2015; 146(1): 100–12.

57. Kruger P, Saffarzadeh M, Weber AN, et al. Neutrophils: Between host defence, immune modulation, and tissue injury. PLoS pathogens 2015; 11(3): e1004651.

58. Tuomanen EI, Saukkonen K, Sande S, Cioffe C, Wright SD. Reduction of inflammation, tissue damage, and mortality in bacterial meningitis in rabbits treated with monoclonal antibodies against adhesion-promoting receptors of leukocytes. The Journal of experimental medicine 1989; 170(3): 959–69.

59. Lowe DM, Demaret J, Bangani N, et al. Differential Effect of Viable Versus Necrotic Neutrophils on Mycobacterium tuberculosis Growth and Cytokine Induction in Whole Blood. Front Immunol 2018; 9: 903.

60. Liu R, Lauridsen HM, Amezquita RA, et al. IL-17 Promotes Neutrophil-Mediated Immunity by Activating Microvascular Pericytes and Not Endothelium. J Immunol 2016; 197(6): 2400–8.

61. Brouwer MC, McIntyre P, Prasad K, van de Beek D. Corticosteroids for acute bacterial meningitis. Cochrane Database Syst Rev 2015; (9): CD004405.

62. Murcia RY, Vargas A, Lavoie JP. The Interleukin-17 Induced Activation and Increased Survival of Equine Neutrophils Is Insensitive to Glucocorticoids. PloS one 2016; 11(5): e0154755.

63. Fragoulis GE, Siebert S, McInnes IB. Therapeutic Targeting of IL-17 and IL-23 Cytokines in Immune-Mediated Diseases. Annu Rev Med 2016; 67: 337–53.

64. Mai NT, Dobbs N, Phu NH, et al. A randomised double blind placebo controlled phase 2 trial of adjunctive aspirin for tuberculous meningitis in HIV-uninfected adults. eLife 2018; 7.

65. Bewersdorf JP, Grandgirard D, Koedel U, Leib SL. Novel and preclinical treatment strategies in pneumococcal meningitis. Current opinion in infectious diseases 2018; 31(1): 85–92.

66. Fisher CJ, Jr., Agosti JM, Opal SM, et al. Treatment of septic shock with the tumor necrosis factor receptor:Fc fusion protein. The Soluble TNF Receptor Sepsis Study Group. The New England journal of medicine 1996; 334(26): 1697–702.

67. Ranieri VM, Thompson BT, Barie PS, et al. Drotrecogin alfa (activated) in adults with septic shock. The New England journal of medicine 2012; 366(22): 2055–64.

